# Distinct modulation of I_h_ by synaptic potentiation in excitatory and inhibitory neurons

**DOI:** 10.1101/2024.09.26.615157

**Authors:** Lotte J. Herstel, Corette J. Wierenga

## Abstract

Selective modifications in the expression or function of dendritic ion channels regulate the propagation of synaptic inputs and determine the intrinsic excitability of a neuron. Hyperpolarization-activated cyclic nucleotide-gated (HCN) channels open upon membrane hyperpolarization and conduct a depolarizing inward current (I_h_). HCN channels are enriched in the dendrites of hippocampal pyramidal neurons where they regulate the integration of synaptic inputs. Synaptic plasticity can bidirectionally modify dendritic HCN channels in excitatory neurons depending on the strength of synaptic potentiation. In inhibitory neurons, however, the dendritic expression and modulation of HCN channels is largely unknown. In this study, we systematically compared the modulation of I_h_ by synaptic potentiation in hippocampal CA1 pyramidal neurons and *stratum Radiatum (sRad)* interneurons. I_h_ properties were similar in inhibitory and excitatory neurons and contributed to resting membrane potential and action potential firing. We found that in *sRad* interneurons, HCN channels were downregulated after synaptic plasticity, irrespective of the strength of synaptic potentiation. This suggest differential regulation of I_h_ in excitatory and inhibitory neurons, possibly signifying their distinct role in network activity.

**Significance statement:** Learning reflects a change in the way information is processed in neuronal circuits. This occurs via changes in synaptic connections and via alterations of intrinsic excitability of neurons. Here we examined how synaptic changes affect properties of HCN channels, which are important ion channels for intrinsic excitability. We found that strong synaptic potentiation leads to opposite changes in HCN channels in CA1 pyramidal neurons and *sRad* interneurons. We speculate that this reflects their differential role in the CA1 network. An upregulation of HCN channels in pyramidal neurons results in a decrease in their excitability, which limits overall network excitation. In contrast, *sRad* interneurons show downregulation of I_h_, and therefore an increased excitability after strong synaptic activation, which will strengthen feedforward inhibition and sharpen activity patterns.

## Introduction

The intrinsic excitability and firing properties of a neuron can be adjusted via selective modifications in the expression or function of specific ion channels (Daoudal and Debanne, 2003; Zhang and Linden, 2003; Debanne et al., 2019). Firing properties are mostly determined by voltage-dependent ion channels in the soma, while ion channels within dendrites regulate the spatial and temporal integration of synaptic inputs along the dendrites (Hoffman et al., 1997; Migliore and Shepherd, 2002; Nolan et al., 2004; Magee and Johnston, 2005; Branco and Häusser, 2010; Oz et al., 2022). Plasticity of neuronal excitability critically contributes to learning and adaptation (Zhang and Linden, 2003; Shah et al., 2010; Hengen et al., 2016; Debanne et al., 2019), memory (Losonczy et al., 2008; Makara et al., 2009), social learning (Gao et al., 2017) and fear conditioning (Carzoli et al., 2023).

Contrary to most other voltage-gated ion channels, hyperpolarization-activated cyclic nucleotide-gated (HCN) channels open when the membrane potential is hyperpolarized. HCN channels are permeable to potassium and sodium ions, which means that they conduct a depolarizing inward current (I_h_). A substantial fraction of HCN channels is open at rest, resulting in a small depolarization of the resting membrane potential (V_rest_). The depolarizing I_h_ increases upon hyperpolarization, and decreases when the membrane gets depolarized. Therefore I_h_ acts to dampen synaptic inputs from both excitatory and inhibitory synapses (Magee, 1999). Because of these distinctive properties, HCN channels are an important contributor to network oscillations (Hu et al., 2009; Zemankovics et al., 2010; Gastrein et al., 2011; Vaidya and Johnston, 2013; Binini et al., 2021). I_h_ also influences the threshold for potentiation of synapses (Losonczy et al., 2008; Shah et al., 2010; Carzoli et al., 2023).

HCN channels are abundantly present in the dendrites of excitatory pyramidal neurons, following a gradient with higher density in the distal dendrites (Lörincz et al., 2002; Harnett et al., 2015). The presence of HCN channels in dendrites reduces dendritic excitability (Magee, 1998; Poolos et al., 2002; Campanac et al., 2008) and their specific distribution along the dendrites results in compartment-specific effects of I_h_ (Harnett et al., 2015; Mäki-Marttunen and Mäki-Marttunen, 2022). Computational modeling showed that the degree of temporal summation of dendritic inputs is primarily determined by the total number of HCN channels and that local dendritic processing is regulated by their dendritic spatial distribution (Angelo et al., 2007). In the hippocampus, HCN channels can also be found in most GABAergic interneurons (Maccaferri and McBain, 1996; Aponte et al., 2006; Anderson et al., 2011; Sekulić et al., 2015). Differences in subcellular localization and/or properties of HCN channels between inhibitory cell types are shown to affect cell-type specific firing properties (Lupica et al., 2001; Aponte et al., 2006; Elgueta et al., 2015; Sekulić et al., 2020), synaptic integration (Sammari et al., 2022), and differential involvement in network activity (Zemankovics et al., 2010; Anderson et al., 2011).

Remarkably, HCN channel properties are strongly regulated by multiple intracellular pathways (Poolos et al., 2002, 2006; Concepcion et al., 2021). HCN channel trafficking in hippocampal neurons is highly dynamic and membrane insertion can occur within minutes (Noam et al., 2010). Impaired regulation of HCN channels is linked to several brain disorders including epilepsy (Albertson et al., 2013; Difrancesco and Difrancesco, 2015) and Fragile X mental retardation (Brager et al., 2012; Brandalise et al., 2020). Synaptic plasticity can locally modify the expression and properties of dendritic HCN channels in excitatory neurons (Daoudal and Debanne, 2003; Wang et al., 2003; Shah et al., 2010). Several studies have described that the modulation of HCN channels in pyramidal neurons depends on both the amplitude and direction of synaptic plasticity (van Welie et al., 2004; Fan et al., 2005; Brager and Johnston, 2007; Campanac et al., 2008; Gasselin et al., 2017), thereby contributing to both Hebbian and homeostatic regulation of intrinsic excitability (Debanne et al., 2019). Plasticity rules are often different for excitatory and inhibitory neurons, reflecting their different roles within the local network (Kullmann et al., 2012; Debanne et al., 2019). It is currently unknown if modulation of HCN channels is differentially regulated in inhibitory and excitatory neurons.

Here, we systematically compared the properties of I_h_ in hippocampal CA1 pyramidal neurons and *stratum Radiatum (sRad)* interneurons. We quantified changes in I_h_ and intrinsic excitability when HCN channels were blocked, following elevated cAMP levels, and after synaptic potentiation. Synaptic potentiation was induced by pairing brief bursts of evoked synaptic potentials with postsynaptic depolarization (theta burst pairing) in pyramidal cells and interneurons. Properties of I_h_ were found generally similar in excitatory and inhibitory cells. However, while pyramidal cells showed an upregulation of I_h_ after strong synaptic potentiation, we found that HCN channels in *sRad* interneurons were downregulated after synaptic plasticity, irrespective of the strength of the synaptic changes. This suggests that pyramidal cells and interneurons express different mechanisms to modulate I_h_, possibly signifying their different roles in the local network.

## Methods

### Animals

All animal experiments were performed in accordance with the guidelines for the welfare of experimental animals and were approved by the local authorities. Mice were kept in standard cages on 12hr light/12hr day cycle under SPF conditions. For this study, GAD65-GFP mice (López-Bendito et al., 2004), bred as a heterozygous line with C57BL/6JRj background, or their wild-type litter mates, of both sexes, were used. In the hippocampus of GAD65-GFP mice, approximately 20% of GABAergic interneurons express GFP. GFP-labelled neurons are mainly Reelin and vasoactive intestinal peptide (VIP)-positive, while they mostly do not express parvalbumin or somatostatin (Wierenga et al., 2010). These cells also express neuropeptide Y, cholecystokinin, calbindin or calretinin. Previous studies have shown that, in the mouse hippocampus, I_h_ could be recorded in most of these interneuron subtypes (Tricoire et al., 2011; Francavilla et al., 2018).

### Slice preparation

Organotypic hippocampal slice cultures were prepared at postnatal day (P)6-8, as previously described by (Stoppini et al., 1991), with some modifications of the protocol. After decapitation the brain was quickly removed and placed in ice cold Gey’s Balanced Salt Solution (GBSS, containing (in mM): 137 NaCl, 5 KCl, 1.5 CaCl_2_, 1 MgCl_2_, 0.3 MgSO_4_, 0.2 KH_2_PO_4_, 0.85 Na_2_HPO_4_) supplemented with 25 mM glucose, 12.5 mM HEPES and 1 mM kynurenic acid, with pH 7.2 and osmolarity ∼320 mOsm/l. Both hippocampi were dissected out and transverse hippocampal slices of 400 μm thick were chopped. The entorhinal cortex (EC) was partially left intact because this area is critical for the development and maintenance of the distal dendritic enrichment of HCN channels in CA1 pyramidal neurons (Shin and Chetkovich, 2007). Slices were placed on Millicell membrane inserts (Millipore, #PICM0RG50) in six-well plates containing 1 ml culture medium (consisting of 48% MEM, 25% HBSS, 25% horse serum, 25 mM glucose, and 12.5 mM HEPES, with pH 7.3 – 7.4 and osmolarity ∼325 mOsm/l) per well. Slice cultures were stored in an incubator (35°C with 5% CO_2_) and the medium was replaced three times a week. Experiments were performed after 10-22 days *in vitro* (DIV).

### Electrophysiology

Before the start of the experiment, a slice was transferred to the recording chamber of the microscope. Artificial cerebral spinal fluid (ACSF; consisting of (in mM): 126 NaCl, 3 KCl, 2.5 CaCl_2_, 1.3 MgCl_2_, 26 NaHCO_3_, 1.25 NaH_2_PO_4_, 20 D-glucose and 1 Trolox, with an osmolarity of 315±10 mOsm/l) was carbonated (95% O_2_, 5% CO_2_), warmed to 30-32 °C and continuously perfused at a speed of ∼1 ml/min. A 4x air objective (Nikon Plan Apochromat) was used to locate the hippocampal CA1 region and cells were visualized with a 60x 1.0 NA water immersion objective (Nikon NIR Apochromat). Whole-cell patch clamp recordings were made of CA1 pyramidal neurons and GFP-expressing interneurons. GFP-positive inhibitory cells were identified in the *statum radiatum*, 100-250 μm from the CA1 pyramidal cell layer, using two-photon fluorescence microscopy. Recording pipettes (resistance of 4-6 MΩ) were filled with an internal solution for measuring excitatory postsynaptic currents (EPSCs; in mM: 140 K-gluconate, 4 KCl, 0.5 EGTA, 10 HEPES, 4 MgATP, 0.4 NaGTP, 4 Na_2_-Phosphocreatine; with pH 7.3 and osmolarity 295±5 mOsm/l) or high chloride internal solution to measure inhibitory postsynaptic currents (IPSCs; in mM: 70 K-gluconate, 70 KCl, 0.5 EGTA, 10 HEPES, 4 MgATP, 0.4 NaGTP, 4 Na_2_-Phosphocreatine; with pH 7.3 and osmolarity 295±5 mOsm/l). Inhibitory currents were isolated by addition of DL-AP5 (50 μM, Tocris) and DNQX (20 μM, Tocris) to the ACSF. Spontaneous action potential firing was prevented by adding 1 μM TTX (Abcam) for the recording of miniature IPSCs. For wash-in experiments, ACSF was substituted with 10 µM ZD7288 (ZD, Sigma) or 25 µM forskolin (FSK, Abcam). During all experiments, cells were kept at a holding potential of -60 mV in both voltage and current clamp. Recordings were excluded when the initial resting membrane potential was above -50 mV. Only interneurons with a V_sag_ larger than 5 mV at -300 pA current injection were included (this cut off was empirically chosen; V_sag_ at -400 pA was > 5 mV for all pyramidal cells). Recordings were acquired using a Multiclamp 700B amplifier (Molecular Devices) with pClamp 10 software.

### Electrical stimulation

To visualize the dendritic arbor after patching, 30 µM Alexa 568 was added to the internal solution (Thermo Fisher Scientific). A concentric bipolar stimulation electrode was placed in a glass pipette filled with ACSF and located in the *stratum radiatum* approximately 100-150 μm from the soma of the patched pyramidal neuron. For interneurons, we placed the stimulation pipette in close proximity to a dendrite 50-100 μm from the soma of the patched cell. For stimulation experiments in excitatory neurons, the CA1 area was surgically isolated to prevent recurrent activity, by a cut separating the CA1 and CA3 region and a cut between the CA1 and the EC. TBS was considered successful if synaptic responses were enhanced right after the stimulation. Synaptic potentiation was determined as the average responses 30 or 60 minutes after TBS, normalized to the average response before stimulation (%EPSC). We discriminated between moderate (<150%) and strong (>150%) potentiation (Campanac et al., 2008). Synaptic responses in inhibitory neurons were consistent over time, but extracellular stimulation in pyramidal neurons often evoked variable and multisynaptic responses that could not be avoided by varying the electrode location or stimulation strength. This may reflect an increased connectivity in slice cultures compared to acute slices. In many pyramidal cells the large variability in the responses made it impossible to unambiguously determine the synaptic potentiation strength after TBS. Experiments in which the extracellular electrode directly stimulated the recorded neuron were disregarded.

For baseline recordings, the stimulus intensity and duration was adjusted to evoke subthreshold postsynaptic currents (evoked PSCs) recorded at 0.1 Hz. We did not block inhibitory currents in these experiments, as washing in GABA_A_ receptor antagonists (bicuculline or gabazine) resulted in massive excitatory currents and cells escaping voltage clamp already at low stimulus strengths, making it impossible to record subthreshold evoked PSCs. Synaptic potentiation was induced after a 10-minute stable baseline with a theta burst stimulation (TBS) paired with postsynaptic depolarization. Theta-modulated burst firing is a behaviorally relevant activity pattern and more efficient in inducing long-term potentiation than other stimulation protocols (Larson and Munkácsy, 2015). One episode of TBS contained 10 bursts at 5 Hz (200 ms intervals) with each burst consisting of five pulses at 100 Hz. One to three TBS episodes were given at 10s intervals. Evoked PSCs were paired with backpropagating action potentials elicited by direct somatic current injection (2 ms, 1 nA; with a 5 ms delay). We included 3 pyramidal cells in which TBS was not paired with postsynaptic depolarization. We continued to record evoked PSCs for at least 30 minutes after the TBS protocol, every two minutes we evoked three responses with an interval of 10s. In between the evoked PSC recordings, we acquired current stimulation recordings.

### Immunochemistry and confocal imaging

Organotypic hippocampal slices were stained for HCN1 to assess the dendritic HCN channel distribution. Slices were fixed in 4% paraformaldehyde for 30 min at room temperature. Next, slices were permeabilized with 0.5% Triton X-100 for 15 min and incubated for 1 h in blocking solution (0.2% Triton X-100 and 10% goat serum). Primary antibodies mouse α-HCN1 (1:1000; NeuroMab, N70/28) and chicken α-MAP2 (1:5000; Abcam/Bio Connect, ab5392) in blocking solution were applied for 24 h at 4°C. Following extensive washing, slices were incubated at room-temperature for 3-4 hours with secondary antibody mixture containing Alexa Fluor 568 anti-mouse (1:500; Thermo Fisher Scientific, A11031) and Alexa Fluor 647 anti-chicken (1:500; Thermo Fisher Scientific, A21449). Slices were mounted with Vectashield medium (Vector laboratories). Confocal laser scanning microscopy images of these slices were acquired on a Zeiss LSM-700 system with a Plan-Apochromat 20x 0.8 NA objective. Z-stack images were acquired with a step size of 1 μm at 1.6 pixels/µm and tiled to construct an image of the whole slice.

### Data analysis

Electrophysiological data was analyzed with Clampfit 10.7 software and custom-written MATLAB scripts. From negative current injection recordings, we determined the negative peak as the maximum voltage (V_max_) and the total hyperpolarization reached at the end of the current injection as steady-state voltage (V_ss_), from which the V_sag =_ V_max_ – V_ss_ was determined. To assess the activation of I_h_, we recorded a two-step protocol, with the first step ranging from −50 to −120 mV (with 10 mV intervals), followed by a second step to −60 mV. Amplitudes of the tail currents (I_tail_) were normalized to the maximum I_tail_ and plotted versus the membrane potential. I_h_ activation curves for each cell were fitted with a Boltzmann function: 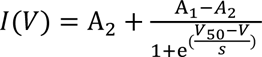, where V is the step voltage, *V_50_* is the half-activation voltage, *s* is the slope factor and A_1_ and A_2_ represent the upper and lower I_tail_ amplitudes. Normalized I_tail_ amplitudes were averaged to show the mean I_h_ activation curve which was refitted by the Boltzmann equation with A_2_ set to 0 and A_1_ set to 1.

To characterize AP firing properties of the recorded neurons, we determined the number of APs for each 500 ms current injection. The number of APs around threshold in Fig. 3G,H,J was determined as the number of APs fired at the smallest current injection step (approximately rheobase). The latency of the first AP was determined at rheobase, except in Fig. 4K, which shows the latency of the first AP for all current steps.

### Statistical analysis

Statistical analysis was performed with Prism 9 (GraphPad). Shapiro-Wilk tests were used to test normality. For normally distributed data points, the statistical significance for paired or unpaired samples was evaluated using the paired or unpaired Student’s t test (t test), respectively. For non-normal distributed unpaired or paired data, we used the nonparametric Mann-Whitney (MW) test or Wilcoxon (W) signed-rank test, respectively. An ANOVA was used to compare normally distributed datasets with multiple measurements with the Friedman test as non-parametric alternative. Kolmogorov-Smirnov (KS) tests were used to compare cumulative distributions. p < 0.05 was considered significant. All data are presented as mean ± SEM.

## Results

### I_h_ contributes to membrane properties in excitatory and inhibitory neurons

In the hippocampal CA1 area, the density of HCN channels on pyramidal neuron dendrites follows a gradient with higher HCN channel density on distal compared to proximal dendrites (Magee, 1998; Lörincz et al., 2002; Harnett et al., 2015). This density gradient of HCN channels efficiently counteracts dendritic filtering (Vaidya and Johnston, 2013) and depends on synaptic activity from entorhinal inputs (Shin and Chetkovich, 2007). Of the four HCN isoforms, HCN1 is highly enriched in the hippocampus (Monteggia et al., 2000; Dougherty et al., 2013). By using antibody staining for HCN1, we confirmed that the HCN channel density gradient is maintained in our organotypic slice cultures (Fig 1A,B).

**Figure 1.**
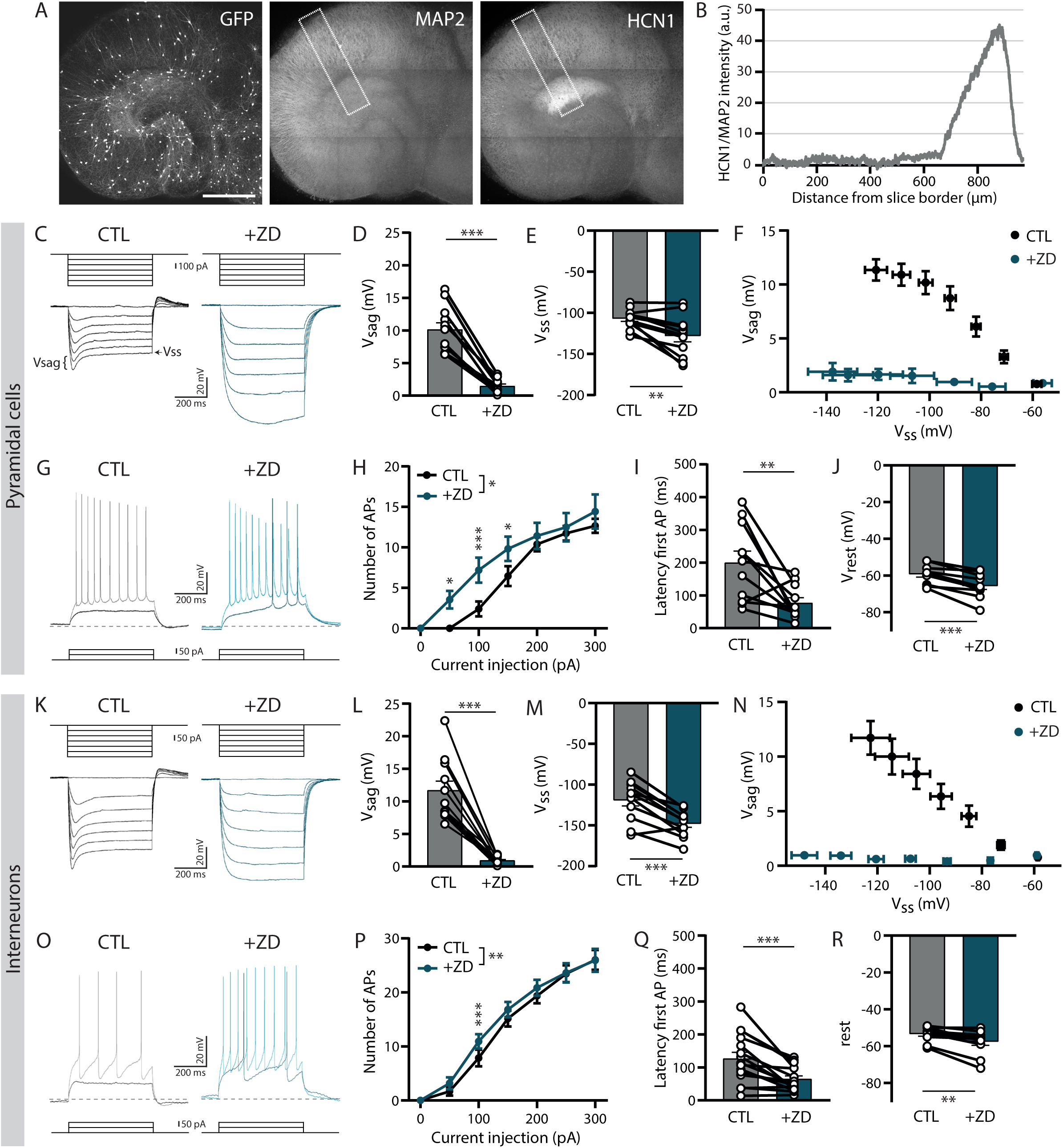
Membrane properties of excitatory and inhibitory neurons are altered by I_h_ blockade. A. Immunostaining for HCN1 and MAP2 in an organotypic hippocampal slice from a GAD65-GFP mouse at DIV 18. Similar antibody labelling was observed in five slices of two mice. Scale bar 500 µm. B. Quantification of the immunostaining in A. HCN1 normalized to MAP2 intensity, plotted over the distance from the slice border in the area indicated by the rectangle in A. C. I_h_ recordings in a pyramidal cell with current injections from 0 to -600 pA (with steps of 100 pA) during baseline (CTL; black) and after wash-in of ZD7288 (+ZD; blue). Voltage sag (V_sag_) and steady-state voltage (V_ss_) are indicated. D. Voltage sag (V_sag_) measured in pyramidal cells at -400 pA current injection (n = 11, p < 00001, t test). E. Steady-state voltage (V_ss_) measured in pyramidal cells at -400 pA current injection (n = 11, p = 0.005, t test). F. Relationship between voltage sag (V_sag_) and steady-state voltage (V_ss_) measured in pyramidal neurons (n = 11, V_sag_: p < 0.0001, V_ss_: p = 0.2 two-way ANOVA). G. Action potential recordings in a pyramidal cell for 50 pA (CTL: black, +ZD: blue) and 100 pA (CTL: grey, +ZD: light blue) current injections. Dotted line indicates the holding potential -60 mV. H. Average number of action potentials (APs) fired in pyramidal neurons for all current injections (n = 11, p = 0.048 two-way ANOVA with multiple comparisons at 50 pA: p = 0.02, 100 pA: p = 0.0006 and 150 pA: p = 0.03). I. Latency of the first AP in pyramidal neurons at the smallest current injection in CTL (n = 11, p = 0.008, t test). J. Resting membrane potential (V_rest_) in pyramidal neurons (n = 11, p < 0.0001, t test). K. I_h_ recordings in an interneuron for current injections from 0 to -300 pA (with steps of 50 pA) during baseline (CTL; black) and wash-in of ZD7288 (+ZD; blue). L. Voltage sag (V_sag_) measured in interneurons at -300 pA current injection (n = 11, p < 0.0001, t test). M. Steady-state voltage (V_ss_) measured in interneurons at -300 pA current injection (n = 11, p = 0.0001, t test). N. Relationship between voltage sag (V_sag_) and steady-state voltage (V_ss_) in interneurons (n = 11, V_sag_: p < 0.0001, V_ss_: p < 0.0001, two-way ANOVA). O. Action potential recordings in an interneuron for 50 pA (CTL: black, +ZD: blue) and 100 pA (CTL: grey, +ZD: light blue) current injections. Dotted line indicates the holding potential -60 mV. P. Average number of action potentials (APs) fired for current injections from 0 to 300 pA in interneurons (n = 15, p = 0.009, two-way ANOVA). Q. Latency of the first AP in interneurons at the smallest current injection in CTL (n = 15, p = 0.0005, t test). R. Resting membrane potential (V_rest_) in interneurons (n = 15, p = 0.002, t test).

We performed whole-cell patch clamp recordings in CA1 pyramidal cells and observed a clear activating I_h_ upon negative current injections in all cells, resulting in the so-called voltage sag (V_sag_) (Fig 1C). The V_sag_ was completely abolished when we blocked HCN channels using the selective blocker ZD7288 (ZD; (Gasparini and DiFrancesco, 1997) (Fig 1D). ZD also resulted in an increase of the steady-state voltage (V_ss_) that was reached during the current injections (Fig 1E). The decrease in V_sag_ and increase in V_ss_ due to the loss of I_h_ were directly related (Fig 1F). When I_h_ was blocked, we also observed an increase in intrinsic excitability in pyramidal neurons, determined by action potential (AP) firing (Fig 1G). The increase in the number of APs was specific for smaller current injections (Fig 1H). Consistently, the latency of the first AP was also decreased upon ZD application (Fig 1I). In addition, we observed a significant hyperpolarization of the resting membrane potential (V_rest_) after adding ZD (Fig 1J). These observations were in good agreement with previous reports (Maccaferri and McBain, 1996; Magee, 1999; Poolos et al., 2002; Fan et al., 2005; Tokay et al., 2009). This shows that I_h_ properties of CA1 pyramidal neurons in our slice cultures were similar to acute slices.

Next, we recorded from GFP-labelled *stratum Radiatum (sRad)* interneurons in slices from GAD65-GFP mice. These interneurons are mostly reelin-positive and target pyramidal cell dendrites (Wierenga et al., 2010). We could measure I_h_ in ∼70% of the GFP-expressing *sRad* interneurons (Fig 1K). As in pyramidal cells, blocking HCN channels with ZD, consistently resulted in an elimination of the V_sag_ and an increase in the V_ss_ (Fig 1L-N). Application of ZD also affected AP firing at small current injections in *sRad* interneurons (Fig 1O-Q), although this effect was diluted due to large variability in firing threshold between the interneurons. I_h_ blockade hyperpolarized V_rest_ (Fig 1R). Our results suggest that I_h_ similarly affects membrane properties and AP firing in CA1 pyramidal cells and *sRad* interneurons.

### Increased cAMP levels shift the activation curve of I_h_ in excitatory and inhibitory neurons

We assessed the activation properties of HCN channels in hippocampal pyramidal neurons and *sRad* interneurons by measuring tail currents (I_tail_) after variable voltage steps. We then constructed an activation curve of I_h_ by plotting the amplitude of the I_tail_ for each voltage step (Fig 2A,B). HCN channels are sensitive to cyclic nucleotides, including cyclic adenosine monophosphate (cAMP; (Biel et al., 2009). Cyclic AMP directly binds to HCN channels and shifts the activation curve of I_h_ towards less negative potentials (Dini et al., 2018; Wainger et al., 2001). To determine how HCN channel activation is regulated by cAMP in CA1 pyramidal cells and *sRad* interneurons, we applied forskolin, which rapidly elevates intracellular cAMP levels via activation of adenylyl cyclase. We observed a small depolarizing shift in the I_h_ activation curve during forskolin application in pyramidal cells (Fig 2C). Normalized I_tail_ during forskolin was smaller for larger voltage steps, resulting in a more depolarized V_50_ after cAMP upregulation (Fig 2D). I_h_ activation curves in interneurons appeared shifted to slightly more hyperpolarized values compared to pyramidal cells under baseline conditions (V_50_ pyramidal cells: -79.6 ± 0.6 mV, V_50_ interneurons: -82.8 ± 1.1 mV, p = 0.054^a^ MW test), while forskolin application resulted in a similar depolarizing shift (Fig 2E,F).

**Figure 2.**
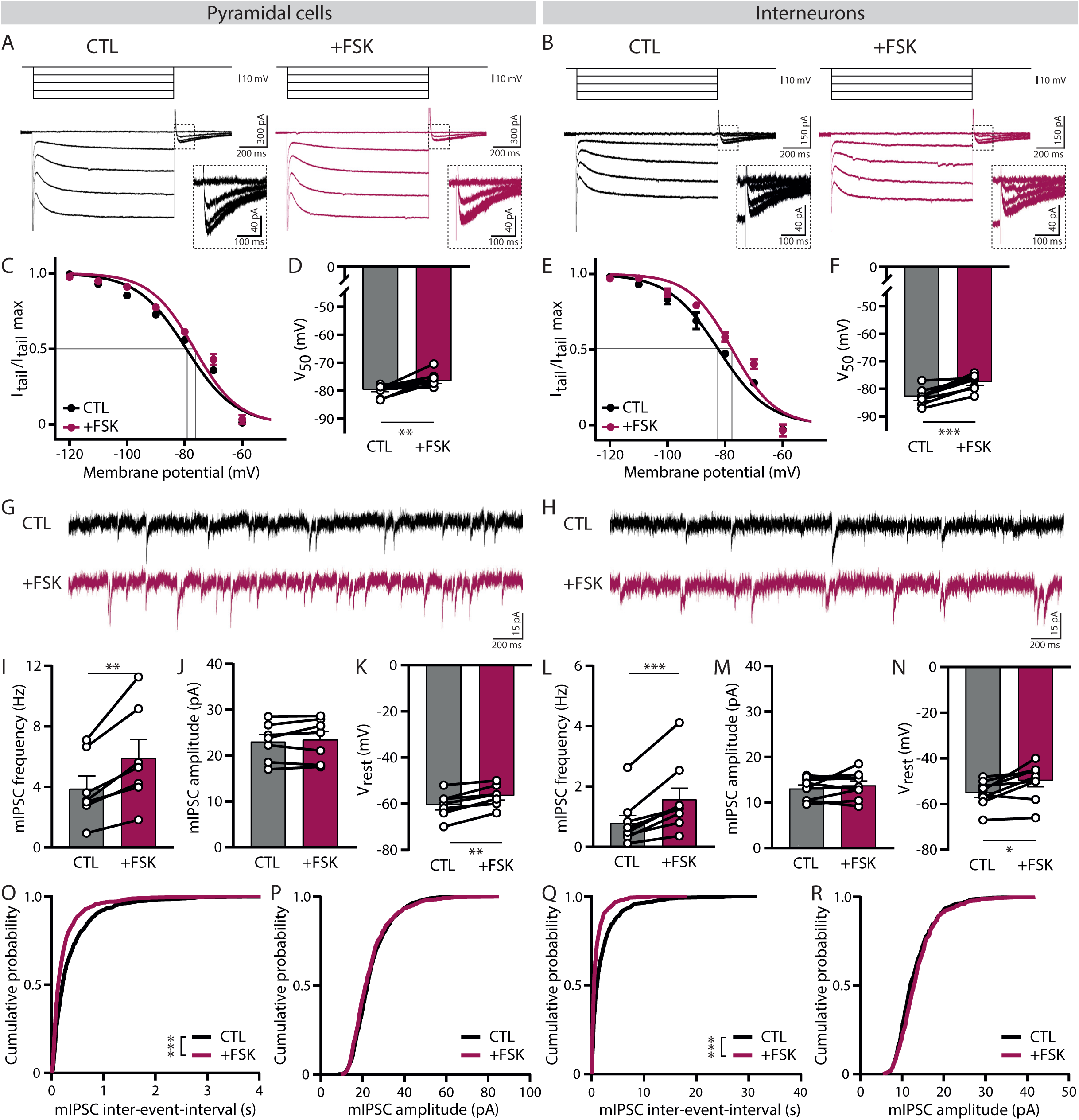
Increase in cAMP with forskolin shifts the activation curve of I_h_ in excitatory and inhibitory neurons. A. I_h_ recordings in a pyramidal cell for voltage steps from -60 to -100 mV (with steps of 10 mV) during baseline (CTL; black) and wash-in of forskolin (+FSK; purple). Insert shows a zoom of the tail currents measured at -60 mV. B. I_h_ recordings in an interneuron for voltage steps from -60 to -100 mV (with steps of 10 mV) during baseline (CTL; black) and wash-in of forskolin (+FSK; purple). Insert shows a zoom of the tail currents measured at -60 mV. C. Average activation curve of I_h_ in pyramidal cells constructed from normalized tail currents (I_tail_/ I_tail_ max) versus membrane potentials (n = 10). D. For each cell activation curves were fitted with a Boltzmann function. V_50_ values for pyramidal cells in CTL and FSK conditions (n = 10, p = 0.01, W test). E. Average activation curve of I_h_ in interneurons constructed from normalized tail currents (I_tail_/ I_tail_ max) versus membrane potentials (n = 8). F. For each cell activation curves were fitted with a Boltzmann function. V_50_ values for interneurons in CTL and FSK conditions (n = 8, p = 0.0004, t test). G. Example of mIPSCs recorded in a pyramidal during baseline (CTL; black) and wash-in of forskolin (+FSK; purple). H. Example of mIPSCs recorded in an interneuron during baseline (CTL; black) and wash-in of forskolin (+FSK; purple). I. Average miniature IPSC frequency in pyramidal cells (n = 7, p = 0.004, t test). J. Average miniature IPSC amplitude in pyramidal cells (n = 7, p = 0.36, t test). K. Resting membrane potential (V_rest_) of pyramidal cells (n = 7, p = 0.002, t test). L. Average miniature IPSC frequency in interneurons (n = 9, p = 0.001, W test). M. Average miniature IPSC amplitude in interneurons (n = 9, p = 0.65, t test). N. Resting membrane potential (V_rest_) of interneurons (n = 9, p = 0.01, t test). O. Cumulative distributions of mIPSC inter-event-interval measured in pyramidal cells (n = 7,p < 0.0001 KS test). For each cell 150 mIPSCs were randomly selected. P. Cumulative distributions of mIPSC amplitude measured in pyramidal cells (n = 7, p = 0.008 KS test). For each cell 150 mIPSCs were randomly selected. Q. Cumulative distributions of mIPSC inter-event-interval measured in interneurons (n = 8, p < 0.0001 KS test). For each cell 75 mIPSCs were randomly selected. R. Cumulative distributions of mIPSCs amplitude measured in interneurons (n = 8, p = 0.007 KS test). For each cell 75 mIPSCs were randomly selected.

We verified that forskolin had fully penetrated the slice by recording miniature inhibitory postsynaptic currents (mIPSCs) (Fig 2G,H). In pyramidal neurons, the frequency of mIPSCs increased almost two-fold within 5 minutes of forskolin wash-in, while mIPSC amplitude was unaffected (Fig 2I,J). This reflects the acute elevation of cAMP levels in presynaptic GABAergic terminals by forskolin (Kaneko and Takahashi, 2004; Huang and Hsu, 2006). In parallel to the increase in mIPSCs, we observed a significant depolarization of V_rest_ in pyramidal cells during forskolin wash-in (Fig 2K), which was consistent with the observed change in the voltage dependence of I_h_ activation. A similar increase in mIPSC frequency and depolarization of V_rest_ was observed in inhibitory neurons (Fig 2L-N). Cumulative distributions of mIPSC amplitudes and interevent intervals were consistent (Fig 2O-R). We noticed that mIPSC frequency remained elevated or at least 30 minutes after we stopped washing in forskolin, while V_rest_ returned to its initial value (data not shown). Together, these data suggest that HCN channels in interneurons and pyramidal cells are comparably sensitive to cAMP and that in both cell types I_h_ gets facilitated by elevating cAMP levels.

### Bidirectional modulation of I_h_ by synaptic potentiation in excitatory neurons

Plasticity of intrinsic excitability via HCN channels has been well described in excitatory neurons (van Welie et al., 2004; Fan et al., 2005; Campanac et al., 2008). To confirm the modulation of I_h_ in CA1 pyramidal neurons in our organotypic slices, we applied TBS paired with postsynaptic depolarization (hereafter referred to as TBS) via an extracellular electrode ∼100-150 µm from the soma to stimulate dendritic synapses (Fig 3A). TBS induced consistent changes in I_h_ upon hyperpolarizing current steps. In some pyramidal cells, V_ss_ was consistently reduced after TBS, indicative of an upregulation of I_h_ (Fig 3B), while in other pyramidal cells V_ss_ was increased after TBS, consistent with a downregulation of I_h_ (Fig 3C). We therefore separated the experiments in two groups, according to the observed change in V_ss_ (Fig 3D). The changes in V_ss_ were accompanied by changes in V_sag_ (Fig 3E-F), in line with either an up- or downregulation of HCN channels after TBS. The change in V_ss_ was well correlated with changes in firing properties around threshold (Fig 3G). An upregulation of I_h_ was accompanied with a small decrease in the number of APs around threshold and an increase in the latency to the first AP (Fig 3H-I). In pyramidal cells in which TBS resulted in a downregulation of I_h_, the number of APs was increased and AP latency was decreased (Fig 3J-K). This is consistent with our earlier observations of the contribution of I_h_ to AP firing (Fig 1H-I). Together these observations indicate that TBS induced HCN channel modulation in pyramidal cells. However, it is important to note here that our data do not exclude additional changes to other ion channels after TBS (Sammari et al., 2022).

**Figure 3.**
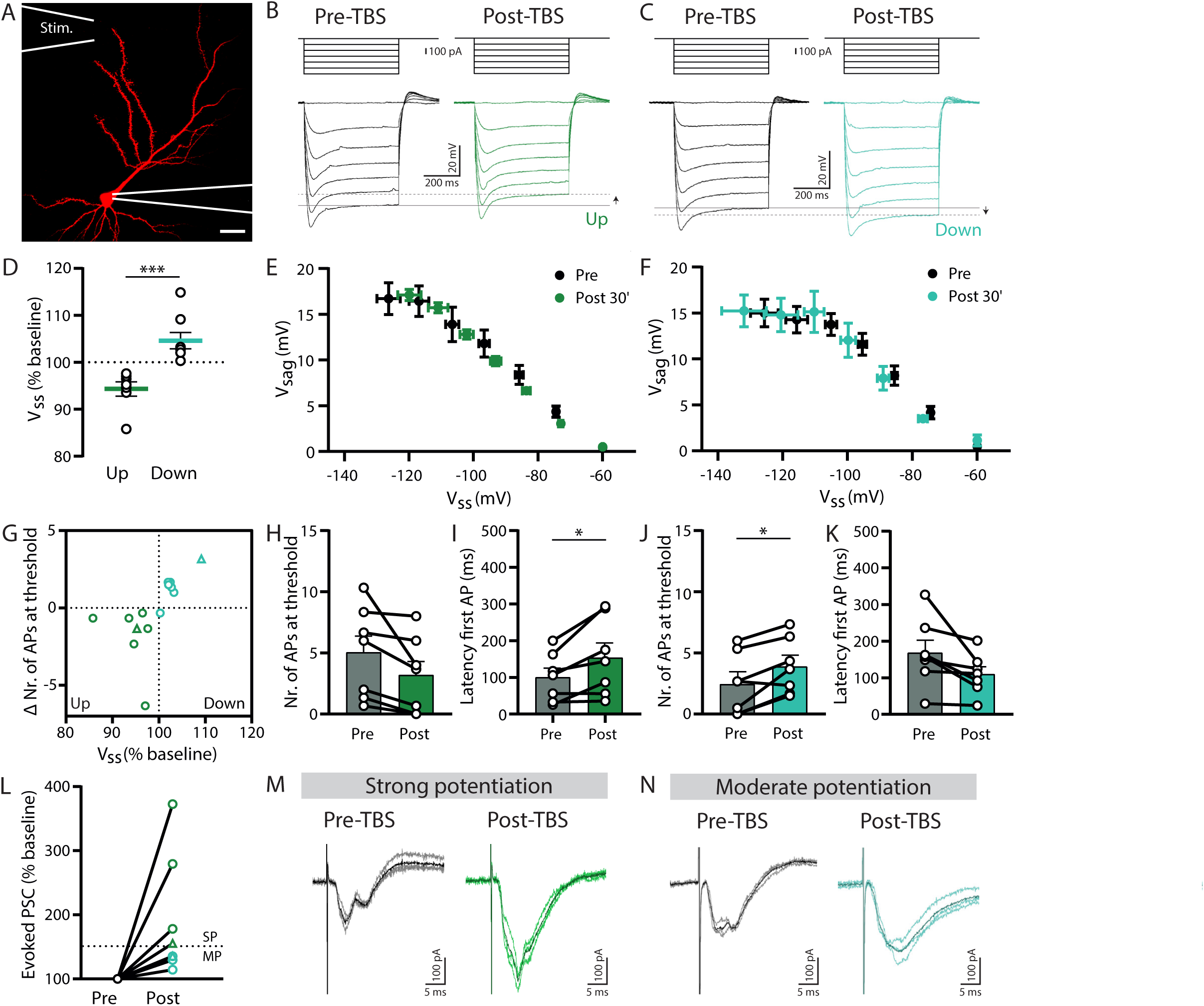
Modulation of I_h_ in pyramidal neurons depends on the strength of synaptic potentiation. A. A pyramidal neuron filled with Alexa 568 via the patch pipette. The location of the stimulation pipette (Stim.) is indicated. Scale bar 20 µm. B. I_h_ recordings for current injections from 0 to -600 pA (with steps of 100 pA) in a pyramidal neuron before (Pre-TBS, black) and after TBS (Post-TBS, green). Arrow indicates an increase in V_ss_, indicating an upregulation of I_h_. C. I_h_ recordings for current injections from 0 to -600 pA (with steps of 100 pA) in a pyramidal neuron before (Pre-TBS, black) and after TBS (Post-TBS, green). Arrow indicates a decrease in V_ss_, indicating a downregulation of I_h_. D. Average change in steady-state voltage (V_ss_, % baseline) at -400 pA current injection for experiments in which I_h_ showed up- and downregulation (Up: n = 7, Down: n = 8, p = 0.0003 MW test). E. Relationship between voltage sag (V_sag_) and steady-state voltage (V_ss_) during baseline (Pre) and 30 minutes post TBS (average of 3 recordings) in pyramidal cells that showed I_h_ upregulation (n = 6, V_sag_: p = 0.04, V_ss_: p < 0.0001, two-way ANOVA). F. Relationship between voltage sag (V_sag_) and steady-state voltage (V_ss_) during baseline (Pre) and 30 minutes post TBS (average of 3 recordings) in pyramidal cells that showed I_h_ downregulation (n = 7, V_sag_: p = 0.8, V_ss_: p = 0.2, two-way ANOVA) G. Correlation between the change in number of APs fired around threshold and the change in steady-state voltage (V_ss_, % baseline). Colors indicate data from pyramidal cells with I_h_ upregulation (dark green) and I_h_ downregulation (light green). Triangles represent the examples shown in M and N. H. Number of APs fired around threshold during baseline (Pre) and 30 minutes post TBS (n = 11, p = 0.056, t test) for pyramidal cells with I_h_ upregulation. I. Latency of the first AP fired during baseline (Pre) and 30 minutes post TBS (n = 11, p = 0.02, t test) for pyramidal cells with I_h_ upregulation. J. Number of APs fired around threshold during baseline (Pre) and 30 minutes post TBS (n = 11, p = 0.011, t test) for pyramidal cells with I_h_ downregulation. K. Latency of the first AP fired during baseline (Pre) and 30 minutes post TBS (n = 11, p = 0.051, t test) for pyramidal cells with I_h_ downregulation. L. Experiments were categorized in moderate synaptic potentiation (MP, evoked PSC <150% of the baseline 20 minutes post-TBS, n = 4) and strong synaptic potentiation (SP, evoked PSC >150%, n = 4). Dashed line indicates 150% potentiation. Triangles represent the examples shown in M and N. M. Example of strong synaptic potentiation. Evoked PSCs during baseline (pre-TBS, average: black, individual traces: grey) and after TBS (post-TBS, average: green, individual traces: light green). N. Example of moderate synaptic potentiation. Evoked PSCs during baseline (pre-TBS, average: black, individual traces: grey) and after TBS (post-TBS, average: green, individual traces: light green).

Previous studies have demonstrated that the direction of HCN channel modulation depends on the strength of synaptic potentiation after TBS. Moderate synaptic potentiation was mostly accompanied by a downregulation of I_h_, while upregulation of I_h_ was triggered after strong synaptic potentiation (Fan et al., 2005; Campanac et al., 2008). In our experiments, the extracellular stimulation often evoked highly varying and multisynaptic responses in many pyramidal cells, which prevented unambiguous quantification of the strength of the synaptic potentiation (also see methods). Although we could not systematically correlate I_h_ changes with synaptic potentiation strength for all experiments, our results were in general agreement with previous studies. Strong synaptic potentiation was observed in pyramidal cells in which I_h_ was upregulated after TBS, while downregulation of I_h_ was associated with more moderate synaptic potentiation (Fig 3L-N). These results indicate a bidirectional modulation of I_h_ in CA1 pyramidal neurons after theta-burst potentiation of synaptic inputs in organotypic hippocampal slices.

### Downregulation of I_h_ by synaptic plasticity in inhibitory neurons independent of the strength of potentiation

Next we recorded from GFP-positive interneurons in the *sRad* of the hippocampal CA1 region (Fig 4A). We only included inhibitory neurons in which I_h_ could be recorded (∼75% of patched cells). We recorded evoked PSCs for 10 minutes (baseline) and then induced synaptic potentiation with TBS paired with postsynaptic depolarization (Fig 4B). Evoked PSCs were recorded for at least 30 minutes after TBS (Fig 4C). Most *sRad* interneurons showed moderate to strong synaptic potentiation and in a few cells (4 out of 11) the extracellular stimulation evoked AP firing after TBS (Fig 4D). We recorded the change in membrane potential for incrementing negative current injections before and after TBS to quantify I_h_ (Fig 4E). We did not observe a significant change in V_sag_ or V_ss_ 30 minutes after TBS, but after 60 minutes the V_ss_ was more hyperpolarized (Fig 4F-H). This was particularly clear in interneurons that had undergone strong synaptic potentiation. All experiments combined, the average V_ss_ at -300 pA current injection was significantly different from baseline after 60 minutes (p = 0.03^b^, one sample t test). We also observed a small increase in AP firing and decrease in latency of the first AP upon small current injections (≤100 pA) 60 minutes after synaptic potentiation (Fig 4I-L). The activation curve of I_h_ did not change after synaptic potentiation when I_tail_ was normalized to their own maximum (Fig 4M-N), indicating that I_h_ activation kinetics were not affected. When we normalized I_tail_ to the maximum I_tail_ in the baseline, the reduction in maximum I_tail_ after synaptic potentiation was clearly noticeable (Fig 4O-P). This shows that TBS-induced synaptic potentiation in *sRad* interneurons results in a downregulation of I_h_ without affecting HCN channel kinetics.

**Figure 4.**
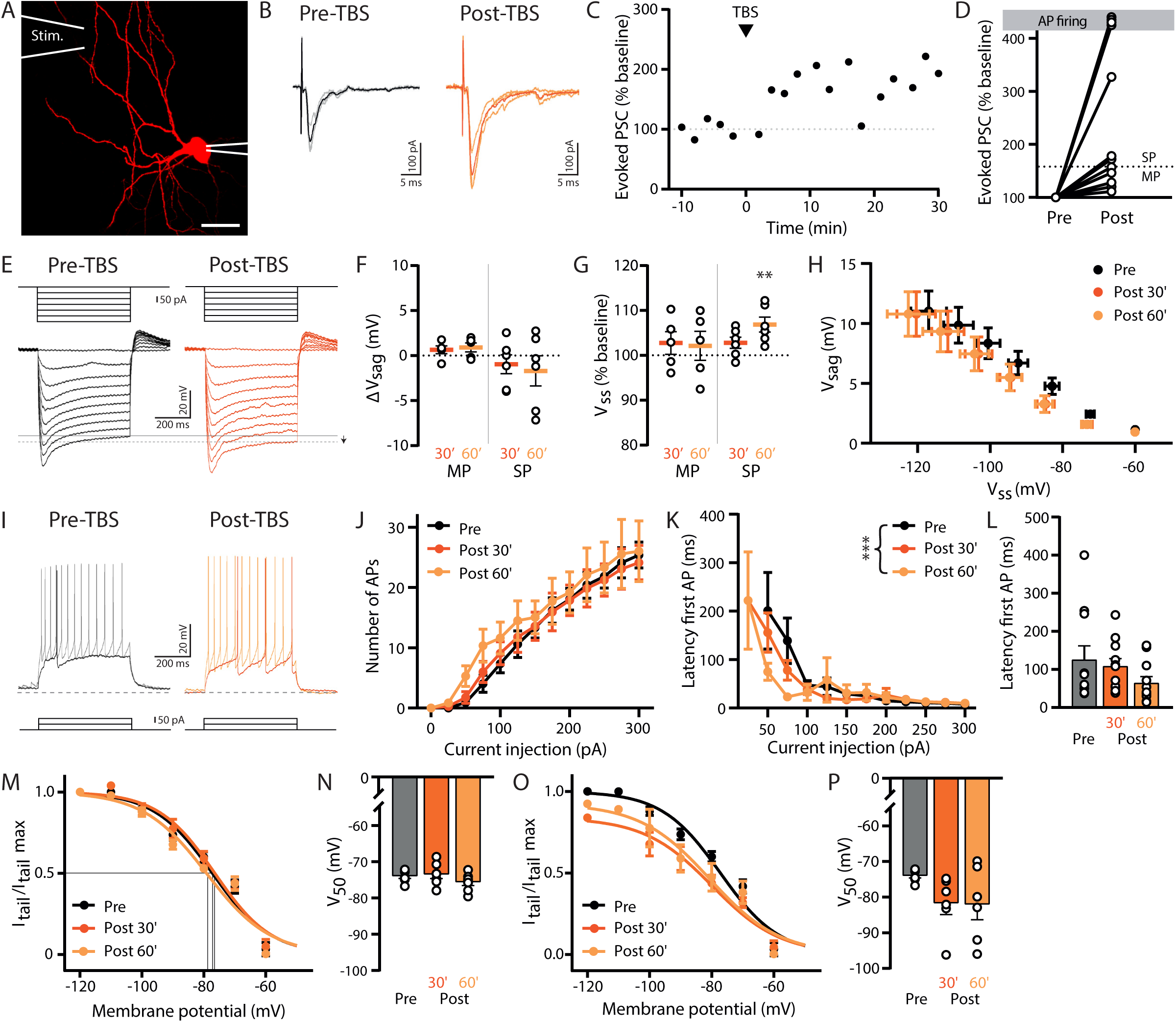
Downregulation of I_h_ in interneurons independent of the strength of synaptic potentiation. A. An interneuron filled with Alexa 568 via the patch pipette. The location of the stimulation (Stim.) pipette is indicated. Scale bar 20 µm. B. Evoked PSCs during baseline (pre-TBS, average: black, individual traces: grey) and after TBS (post-TBS, average: orange, individual traces: light orange). C. Evoked PSC amplitude (% baseline) over time for the experiment shown in B. D. Experiments were categorized in moderate synaptic potentiation (MP, evoked PSC <150% of baseline 0-20 minutes after the TBS, n = 5) and strong synaptic potentiation (SP, evoked PSC >150%, n = 6). Dashed line indicates 150% potentiation. E. I_h_ recordings in an interneuron for current injections from 0 to -300 pA (with steps of 50 pA) before (Pre-TBS, black) and after TBS (Post-TBS, orange). Arrow indicates a decrease in V_ss_, indicating a downregulation of I_h_. F. Voltage sag (ΔV_sag_) at -300 pA current injection 30 and 60 minutes post TBS (average of 3 recordings) for experiments with moderate and strong synaptic potentiation (MP n = 5 and SP n = 6). G. Steady-state voltage (V_ss,_ % baseline) at -300 pA current injection 30 and 60 minutes post TBS (average of 3 recordings) for experiments with moderate and strong synaptic potentiation (MP n = 5 and SP n = 6, SP 60’: p = 0.009, one sample t-test). H. Relationship between voltage sag (V_sag_) and steady-state voltage (V_ss_) during baseline (Pre), 30 and 60 minutes post TBS (average of 3 recordings) in interneurons (n = 11, V_sag_: p = 0.5, V_ss_: p = 0.06, two-way ANOVA). I. Action potential recordings in an interneuron for 50 pA (Pre-TBS: black, Post-TBS: orange) and 100 pA (Pre-TBS: grey, Post-TBS: light orange) current injections. Dotted line indicates -60 mV. J. Average number of action potentials (APs) fired in interneurons for all current injections (n = 11, p = 0.9, two-way ANOVA). K. Latency of the first AP fired in interneurons for all current injections during baseline (Pre), 30 and 60 minutes post-TBS (n = 11, p < 0.0001 mixed effects model). L. Latency of the first AP in interneurons at the smallest current injection Pre-TBS (n = 11, p = 0.1, Friedman test). M. Average I_h_ activation curve constructed by normalizing tail currents to the maximum I_tail_ per condition (n = 7). Curves were fitted with Boltzmann functions. N. V_50_ values determined from individual I_h_ activation curves as shown in N (n = 7, p = 0.3, one-way ANOVA). O. Average I_h_ activation curve constructed by normalizing tail currents (I_tail_) to the maximum I_tail_ during baseline (Pre-TBS, n = 7). Curves were fitted with Boltzmann functions. P. V_50_ values determined from individual I_h_ activation curves as shown in O (n = 7, p = 0.1, one-way ANOVA).

## Discussion

In this study, we compared I_h_ in pyramidal cells and *sRad* interneurons in the hippocampal CA1 area. We found that I_h_ properties were similar and that I_h_ contributes significantly to V_rest_ and AP firing in both types of neurons. TBS induced synaptic potentiation in both cell types, which was associated with either an up- or downregulation of I_h_ in pyramidal cells, while TBS only induced downregulation of I_h_ in *sRad* interneurons. Our data indicate that HCN channel regulation is cell-type specific and we speculate that this may signify the different roles of these neurons in the local network.

The density gradient and recordings of I_h_ (i.e. V_sag_ and V_ss_) in CA1 pyramidal cells in our experiments were similar to previous reports in acute slices (Magee, 1998; Wilkars et al., 2012; Srinivas et al., 2017). In CA1 pyramidal cells, dendritic HCN channels are distributed following a density gradient with higher densities further away from the soma (Lörincz et al., 2002; Harnett et al., 2015). The cellular mechanism underlying the dendritic gradient of I_h_ remains unclear (Kupferman et al., 2014; Meseke et al., 2018), but it was shown to require intact projections from the entorhinal cortex (Shin and Chetkovich, 2007). We keep a part of the entorhinal cortex attached to our hippocampal slices to maintain the CA1 network architecture (Brewster et al., 2006; Shin and Chetkovich, 2007). After two weeks in culture, our hippocampal slices would be roughly equivalent to the 3^rd^ postnatal week *in vivo* (De Simoni et al., 2003). It is less clear if HCN channels also have a specific cellular distribution in interneurons. In parvalbumin cells, HCN channels are highly enriched in axons (Aponte et al., 2006; Elgueta et al., 2015; Roth and Hu, 2020), but in other interneurons HCN channels appear to be localized mostly in the soma and proximal dendrites (Anderson et al., 2011; Sekulić et al., 2015, 2020).

From our experiments, there are no indications that the properties of HCN channels are different in inhibitory and excitatory neurons. We found that I_h_ affects passive membrane properties, V_rest_ and input resistance similarly in hippocampal CA1 pyramidal cells and *sRad* interneurons. I_h_ dampens intrinsic excitability, and blocking HCN channels with ZD led to a small increase in AP firing and decrease in V_rest_. Forskolin application had two independent effects: it triggered an increase in mIPSC frequency and shifted the I_h_ activation curve in both cell types. Forskolin activates the enzyme adenylyl cyclase, which converts ATP to the second messenger cAMP. Cyclic AMP is known to stimulate presynaptic vesicle release via activation of protein kinases (Weisskopf et al., 1994; Nicoll and Schmitz, 2005; Antoni, 2012). The forskolin-mediated increase in mIPSC frequency was similar in pyramidal cells and interneurons, and likely reflects a general increase in vesicle release in presynaptic inhibitory terminals in the brain slice (Kaneko and Takahashi, 2004; Huang and Hsu, 2006). In parallel, we observed a depolarizing shift in I_h_ activation in both cell types, consistent with previous reports that cAMP can directly influence the gating properties of HCN channels (Wainger et al., 2001; Magee et al., 2015; Dini et al., 2018). Due to the depolarizing shift in gating, more HCN channels are open at rest resulting in a small, but significant, depolarization of V_rest_ (Gambardella et al., 2012). We noticed that the V_rest_ depolarization was abolished when forskolin application ended, while mIPSC frequency remained elevated. This is consistent with the notion that mIPSC increase is mediated via cAMP-dependent kinases (Capogna et al., 1995; Fernandes et al., 2015), while the effect on HCN channels and V_rest_ is directly mediated by cAMP (Lüthi and McCormick, 1999).

The most important observation in this study was that synaptic potentiation had a differential effect on HCN channels in pyramidal cells and *sRad* interneurons. The changes in I_h_ were clear from changes in V_sag_ and V_ss_ during hyperpolarizing current steps. However, it is important to note here that we did not perform pharmacological controls and we therefore cannot exclude additional changes to other ion channels after TBS (Sammari et al., 2022). Previous studies showed that modulation of HCN channels in hippocampal pyramidal neurons depends on the strength of synaptic potentiation. Large synaptic potentiation causes an upregulation of I_h_ and therefore a decrease in intrinsic excitability, whereas moderate potentiation results in a downregulation of I_h_ associated with increased intrinsic excitability (van Welie et al., 2004; Fan et al., 2005; Campanac et al., 2008; Debanne et al., 2019). We used TBS to induce synaptic potentiation in pyramidal cells and interneurons. We observed an increase in I_h_ only in some pyramidal cells, while in others I_h_ was downregulated after TBS. Although we could not directly relate this to the strength of synaptic potentiation in our experiments, our results clearly demonstrate bidirectional I_h_ modulation in CA1 pyramidal cells, in line with these previous studies. In contrast, we always observed a reduction in I_h_ in *sRad* interneurons after synaptic potentiation. The difference in I_h_ modulation between pyramidal cells and interneurons cannot be explained by a reduced efficiency of synaptic potentiation in the *sRad* interneurons. In fact, synaptic stimulation resulted in AP firing in some interneurons after TBS, indicating that maximal synaptic potentiation was reached. Our data suggest that inhibitory neurons only exhibit a mechanism to downregulate HCN channels and lack a mechanism for I_h_ upregulation.

Previous studies have identified several molecular mechanisms that regulate surface expression and activation properties of HCN channels. The gating properties of I_h_ are modulated by cAMP and cGMP (Wainger et al., 2001; Maroso et al., 2016), but also by PIP_2_ (Pian et al., 2006; Zolles et al., 2006). In addition, HCN proteins contain multiple phosphorylation sites by which the number and properties of functional HCN channels in the membrane are regulated (Williams et al., 2015; Concepcion et al., 2021). TBS can increase intracellular cAMP levels (Nguyen and Kandel, 1997; Lüthi and McCormick, 1999) and an increase in cAMP enhances I_h_ (Lüthi and McCormick, 1999). We found that HCN channels in excitatory and inhibitory cells are equally sensitive to changes in cAMP levels upon forskolin application. The synaptic potentiation-driven reduction in I_h_ conductance in interneurons was not accompanied by a change in activation kinetics (Fig 4M-P), which suggests that TBS did not elevate cAMP levels in these interneurons. Previous studies have reported downregulation of I_h_ also in excitatory neurons, when synaptic potentiation after TBS was moderate. In that case, the reduction in I_h_ conductance also occurred without a change in the activation curve (Campanac et al., 2008), very similar to what we observe in *sRad* interneurons. This suggests that an increase in cAMP may be required for I_h_ upregulation after strong synaptic potentiation in pyramidal cells, but that moderate synaptic potentiation does not affect cAMP levels. The observed difference in I_h_ modulation after TBS stimulation in pyramidal cells and interneurons may therefore suggest a differential effect of the synaptic stimulation on intracellular cAMP levels. In addition, baseline cAMP levels in CA1 pyramidal cells and *sRad* interneurons may already be different as suggested by the apparent difference in V_50_ values of the I_h_ activation curve in these cells (Fig 2C and 2E). Future studies could employ novel PKA sensors (Ma et al., 2018) to directly compare cAMP dynamics in excitatory and inhibitory cells after synaptic potentiation. In addition to cAMP, upregulation of I_h_ after synaptic potentiation may involve CaMKII and NMDA receptor activation (van Welie et al., 2004; Fan et al., 2005). It is also possible that the absence of I_h_ upregulation in *sRad* interneurons is due to the cell-type specific expression of some molecular components of the upregulation pathway (e.g. expression of CaMKIIα is much lower in interneurons; Liu and Jones, 1996; Sík et al., 1998; Keaveney et al., 2020; Veres et al., 2023).

Downregulation of I_h_ can be mediated via PKC activation (Brager and Johnston, 2007; Williams et al., 2015) or via PLC-mediated depletion of PIP_2_ (Pian et al., 2006; Zolles et al., 2006). A recent report described that downregulation of HCN channels after synaptic potentiation in OLM cells is mediated by mGluR1 activation (Sammari et al., 2022). The downregulation of I_h_ and increase in AP firing that we observed here are very similar to what was reported in OLM cells and it is therefore tempting to speculate that a similar pathway is involved in *sRad* interneurons. We noticed that the increase in V_ss_ in *sRad* interneurons became significant only after 60 minutes, suggesting that I_h_ downregulation is slower in inhibitory neurons compared to excitatory neurons (Campanac et al., 2008) and OLM cells (Sammari et al., 2022).

Intrinsic excitability and firing properties are highly cell-type specific, reflecting specific genetic programs within cell types, which will be influenced by network activity patterns. In addition, cell-type specific regulation of ion channels may depend on the role of the neuron in the network. Within neuronal networks, plasticity of excitation is usually accompanied by plasticity of inhibition to ensure fidelity of information processing and to enable computational flexibility (Carvalho and Buonomano, 2009; Froemke, 2015; Herstel and Wierenga, 2021). For both excitation and inhibition, plasticity of synaptic connections and intrinsic excitability are coordinated to achieve changes in network function (Kullmann et al., 2012; Gao et al., 2017; Debanne et al., 2019). Activity- and context-dependent recruitment of inhibitory cells is important for information processing in neuronal networks, and behavioral flexibility. It is therefore not surprising that different plasticity rules apply for feedforward and feedback inhibition (Lamsa et al., 2005, 2007; Sambandan et al., 2010), reflecting their different role in the network. The GFP-expressing *sRad* interneurons in slices from GAD65-GFP mice consist of a broad population of different interneurons subtypes, but they mostly target the dendrites of CA1 pyramidal cells (Wierenga et al., 2010) and provide feedforward inhibition to CA1 pyramidal cells (Cope et al., 2002; Wierenga and Wadman, 2003; Lamsa et al., 2005; Milstein et al., 2015). The upregulation of I_h_ in pyramidal cells after strong synaptic potentiation is thought to restrict uncontrolled activity (Debanne et al., 2019). Here we found that *sRad* interneurons respond to synaptic plasticity by decreasing I_h_, which will increase their excitability. An increase in interneuron excitability after strong network activity makes sense from a network point of view. We speculate that the increase in excitability of *sRad* interneurons is important to strengthen feedforward inhibition and helps to sharpen activity patterns (Lamsa et al., 2005; Kullmann et al., 2012). Plasticity of HCN channels provides an intracellular mechanism to adjust and finely regulate neuronal excitability in reaction to synaptic stimulation. Our data highlight that regulation mechanisms for HCN channels vary between neuronal cell types.

## Supporting information

statistical table

