## Supplementary material for "Distinct modulation of I_h_ by synaptic potentiation in excitatory and inhibitory neurons": statistical table

| Fig. | Data structure | Type of test | Sample size (n) | P-value | Power |
|---|---|---|---|---|---|
| 1D | Normal distribution | Paired Student's *t* test | 11 cells, 11 slices, 8 mice | p < 00001 | Mean diff: -8.7 ± 1.0, 95% CI: -11.0 to -6.4 |
| 1E | Normal distribution | Paired Student's *t* test | 11 cells, 11 slices, 8 mice | p = 0.005 | Mean diff: -20.3 ± 5.6, 95% CI: -32.7 to -7.9 |
| 1F $V_{sag}$ | Normal distribution | Mixed effects model<br>*Current x Condition*<br>Šídák's multiple comparisons test | 11 cells, 11 slices, 8 mice | F(6,68) = 10.45<br>p < 0.0001 | -200 pA: Mean diff: 6.1 ± 1.1, 95% CI: 3.1 to 9.1<br>-300 pA: Mean diff: 7.2 ± 1.1, 95% CI: 4.2 to 10.2<br>-400 pA: Mean diff: 8.3 ± 1.1, 95% CI: 5.232 to 11.28<br>-500 pA: Mean diff: 9.3 ± 1.1, 95% CI: 6.2 to 12.4<br>-600 pA: Mean diff: 9.3 ± 1.1, 95% CI: 6.2 to 12.4 |
| 1F $V_{ss}$ | Normal distribution | Two way repeated measures ANOVA<br>*Current x Condition* | 11 cells, 11 slices, 8 mice | F(6,70) = 1.35<br>p = 0.25 | |
| 1H | Normal distribution | Two way repeated measures ANOVA<br>*Current x Condition*<br>Šídák's multiple comparisons test | 11 cells, 11 slices, 8 mice | F(6,66) = 2.26<br>p = 0.048 | 50 pA: Mean diff: -3.5 ± 1.1, 95% CI: -6.6 to -0.3, p = 0.02<br>100 pA; Mean diff: -4.8 ± 1.1, 95% CI: -8.0 to -1.6, p = 0.0006<br>150 pA: Mean diff: -3.4 ± 1.1, 95% CI: -6.5 to -0.2, p = 0.03 |
| 1I | Normal distribution | Paired Student's *t* test | 11 cells, 11 slices, 8 mice | p = 0.008 | Mean diff.: -122.8 ± 36.8, 95% CI: -204.9 to -40.7 |
| 1J | Normal distribution | Paired Student's *t* test | 11 cells, 11 slices, 8 mice | p < 0.0001 | Mean diff.: -6.4 ± 1.0, 95% CI: -8.5 to -4.3 |
| 1L | Normal distribution | Paired Student's *t* test | 11 cells, 11 slices, 9 mice | p < 0.0001 | Mean diff: -10.8 ± 1.4, 95% CI: -13.9 to -7.6 |
| 1M | Normal distribution | Paired Student's *t* test | 11 cells, 11 slices, 9 mice | p = 0.0001 | Mean diff: -28.8 ± 4.8, 95% CI: -39.6 to -18.0 |
| 1N $V_{sag}$ | Normal distribution | Two way repeated measures ANOVA<br>*Current x Condition*<br>Šídák's multiple comparisons test | 11 cells, 11 slices, 9 mice | F(6,70) = 12.19<br>p < 0.0001 | -100 pA: Mean diff: 4.1 ± 1.1, 95% CI: 0.9 to 7.3, p = 0.004<br>-150 pA: Mean diff: 5.7 ± 1.1, 95% CI: 2.5 to 8.9, p < 0.0001<br>-200 pA: Mean diff: 7.8 ± 1.1, 95% CI: 4.6 to 11.0, p < 0.0001<br>-250 pA: Mean diff: 9.1 ± 1.1, 95% CI: 5.9 to 12.2, p < 0.0001<br>-300 pA: Mean diff: 10.8 ± 1.1, 95% CI: 7.6 to 13.9, p < 0.0001 |
| 1N $V_{ss}$ | Normal distribution | Two way repeated measures ANOVA<br>*Current x Condition*<br>Šídák's multiple comparisons test | 11 cells, 11 slices, 9 mice | F(6,70) = 9.39<br>p < 0.0001 | -100 pA: Mean diff: 9.9 ± 3.3, 95% CI: 0.8 to 18.9, p = 0.03<br>-150 pA: Mean diff: 13.8 ± 3.3, 95% CI: 4.7 to 22.9, p < 0.0005<br>-200 pA: Mean diff: 18.0 ± 3.3, 95% CI: 8.9 to 27.1, p < 0.0001<br>-250 pA: Mean diff: 22.6 ± 3.3, 95% CI: 13.5 to 31.7, p < 0.0001<br>-300 pA: Mean diff: 28.8 ± 3.3, 95% CI: 19.7 to 37.9, p < 0.0001 |
| 1P | Normal distribution | Two way repeated measures ANOVA<br>*Current x Condition*<br>Šídák's multiple comparisons test | 15 cells, 15 slices, 11 mice | F(6,98) = 3.17<br>p = 0.007 | 100 pA: Mean diff: -3.1 ± 0.7, 95% CI: -4.9 to -1.3, p < 0.0001 |
| 1Q | Normal distribution | Paired Student's *t* test | 15 cells, 15 slices, 11 mice | p = 0.0005 | Mean diff: -61.9 ± 13.7, 95% CI: -91.2 to -32.6 |
| 1R | Non-normal distribution | Wilcoxon signed-rank test | 11 cells, 11 slices, 8 mice | p = 0.002 | |
| 2D | Non-normal distribution | Wilcoxon signed-rank test | 10 cells, 10 slices, 8 mice | p = 0.01 | |
| 2F | Normal distribution | Paired Student's *t* test | 8 cells, 8 slices, 6 mice | p = 0.0004 | Mean diff: 5.3 ± 0.8, 95% CI: 3.3 to 7.2 |
| 2I | Normal distribution | Paired Student's *t* test | 7 cells, 7 slices, 6 mice | p = 0.004 | Mean diff: 2.0 ± 0.4, 95% CI: 1.0 to 3.1 |
| 2J | Normal distribution | Paired Student's *t* test | 7 cells, 7 slices, 6 mice | p = 0.36 | Mean diff: 0.5 ± 0.5, 95% CI: -0.7 to 1.6 |
| 2K | Normal distribution | Paired Student's *t* test | 7 cells, 7 slices, 6 mice | p = 0.002 | Mean diff: 4.0 ± 0.8, 95% CI: 2.2 to 5.9 |

| | | | | | |
|---|---|---|---|---|---|
| 2L | Non-normal distribution | Wilcoxon signed-rank test | 9 cells, 9 slices, 6 mice | p = 0.004 | |
| 2M | Normal distribution | Paired Student's *t* test | 9 cells, 9 slices, 6 mice | p = 0.35 | Mean diff: 0.7 ± 0.7, 95% CI: -0.9 to 2.3 |
| 2N | Normal distribution | Paired Student's *t* test | 9 cells, 9 slices, 6 mice | p = 0.01 | Mean diff: 5.3 ± 1.7, 95% CI: 1.5 to 9.2 |
| 2O | Normal distribution | Kolmogorov-Smirnov test | 1015 mIPSCs, 7 cells | p < 0.0001 | |
| 2P | Normal distribution | Kolmogorov-Smirnov test | 1015 mIPSCs, 7 cells | p = 0.008 | |
| 2Q | Normal distribution | Kolmogorov-Smirnov test | 675 mIPSCs, 9 cells | p < 0.0001 | |
| 2R | Normal distribution | Kolmogorov-Smirnov test | 675 mIPSCs, 9 cells | p = 0.007 | |
| *a** | Non-normal distribution | Mann Whitney *U* test | | p = 0.054 | |
| 3D | Non-normal distribution | Mann Whitney *U* test | Up: 7 cells, 7 slices, 6 mice<br>Down: 8 cells, 8 slices, 8 mice | p = 0.0003 | |
| 3E<br>$V_{sag}$ | Normal distribution | Two way repeated measures ANOVA<br>*Current x Condition*<br>Šídák's multiple comparisons test | 6 cells, 6 slices, 5 mice | F(6,33) = 2.52<br>p = 0.04 | -100 pA: Mean diff: 1.7 ± 0.5, 95% CI: 0.2 to 3.3, p = 0.02<br>-150 pA: Mean diff: 1.9 ± 0.5, 95% CI: 0.4 to 3.4, p = 0.007 |
| 3F<br>$V_{ss}$ | Normal distribution | Two way repeated measures ANOVA<br>*Current x Condition*<br>Šídák's multiple comparisons test | 6 cells, 6 slices, 5 mice | F(6,33) = 11.89<br>p < 0.0001 | -100 pA: Mean diff: -2.2 ± 0.7, 95% CI: -4.1 to -0.3, p = 0.02<br>-150 pA: Mean diff: -3.6 ± 0.7, 95% CI: -5.5 to -1.7, p < 0.0001<br>-200 pA: Mean diff: -4.5 ± 0.7, 95% CI: -6.4 to -2.6, p < 0.0001<br>-250 pA: Mean diff: -6.0 ± 0.7, 95% CI: -8.0 to -3.9, p < 0.0001<br>-300 pA: Mean diff: -6.4 ± 0.7, 95% CI: -8.5 to -4.4, p < 0.0001 |
| 3F<br>$V_{sag}$ | Normal distribution | Two way repeated measures ANOVA<br>*Current x Condition*<br>Šídák's multiple comparisons test | 7 cells, 7 slices, 7 mice | p = 0.81 | |
| 3F<br>$V_{ss}$ | Normal distribution | Two way repeated measures ANOVA<br>*Current x Condition*<br>Šídák's multiple comparisons test | 7 cells, 7 slices, 7 mice | p = 0.18 | |
| 3H | Normal distribution | Paired Student's *t* test | 7 cells, 7 slices, 6 mice | p = 0.056 | Mean diff: -1.9 ± 0.8, 95% CI: -3.8 to 0.1 |
| 3I | Normal distribution | Paired Student's *t* test | 7 cells, 7 slices, 6 mice | p = 0.02 | Mean diff: 54.0 ± 18.0, 95% CI: 9.9 to 98.1 |
| 3J | Normal distribution | Paired Student's *t* test | 8 cells, 8 slices, 8 mice | p = 0.011 | Mean diff: 1.4 ± 0.4, 95% CI: 0.5 to 2.4 |
| 3K | Normal distribution | Paired Student's *t* test | 8 cells, 8 slices, 8 mice | p = 0.051 | Mean diff: -58.1 ± 23.9, 95% CI: -116.6 to 0.3 |
| 4F | Normal distribution | One sample Student's *t* tests | MP: 5 cells, 5 slices, 4 mice<br>SP: 6 cells, 6 cells, 6 mice | MP 30': p = 0.21<br>SP 30': p = 0.40<br>MP 60': p = 0.14<br>SP 60': p = 0.36 | MP 30': Discr.: 0.6 ± 0.4, 95% CI: -0.6 to 1.8<br>SP 30': Discr.: -1.0 ± 1.0, 95% CI: -3.7 to 1.7<br>MP 60': Discr.: 0.9 ± 0.5, 95% CI: -0.5 to 2.3<br>SP 60': Discr.: -1.7 ± 1.7, 95% CI: -6.0 to 2.6 |
| 4G | Normal distribution | One sample Student's *t* tests | MP: 5 cells, 5 slices, 4 mice<br>SP: 6 cells, 6 cells, 6 mice | MP 30': p = 0.34<br>SP 30': p = 0.08<br>MP 60': p = 0.55<br>SP 60': p = 0.009 | MP 30': Discr.: 2.7 ± 2.5, 95% CI: -4.3 to 9.8<br>SP 30': Discr.: 2.8 ± 1.3, 95% CI: -0.4 to 6.0<br>MP 60': Discr.: 2.119 ± 3.2, 95% CI: -6.9 to 11.1<br>SP 60': Discr.: 6.858 ± 1.7, 95% CI: 2.6 to 11.1 |
| 4H<br>$V_{sag}$ | Normal distribution | Two way repeated measures ANOVA<br>*Current x Condition* | 11 cells, 11 slices, 7 mice | F(12,140) = 0.94<br>p = 0.51 | |
| $V_{ss}$ | Normal distribution | Two way repeated measures ANOVA | 11 cells, 11 slices, 7 mice | F(12,140) = 1.77 | |

| | | | | | |
|---|---|---|---|---|---|
| | | *Current x Condition* | | p = 0.06 | |
| 4J | Normal distribution | Two way repeated measures ANOVA | 11 cells, 11 slices, 7 mice | F(24,208) = 0.33<br>p = 0.99 | |
| 4K | Normal distribution | Mixed effects model<br>*Current x Condition* | 11 cells, 11 slices, 7 mice | F(20,172) = 5.33<br>p < 0.0001 | 50 pA 30' vs 60': Mean diff: 81.9 ± 21.8, 95% CI: 3.9 to 159.2, p = 0.04<br>75 pA pre vs 60': Mean diff: 116.3 ± 35.9, 95% CI: 6.1 to 226.5, p = 0.04<br>125 pA pre vs 30': Mean diff: 24.7 ± 7.5, 95% CI: 1.6 to 47.7, p = 0.04<br>150 pA pre vs. 30': Mean diff: 11.5 ± 3.6, 95% CI: 1.1 to 21.9, p = 0.03 |
| 4L | Non-normal distribution | Friedman test | 11 cells, 11 slices, 7 mice | p = 0.11 | |
| 4N | Normal distribution | One way ANOVA | 7 cells, 7 slices, 5 mice | F(2,18) = 1.49<br>p = 0.25 | |
| 4P | Normal distribution | One way ANOVA | 7 cells, 7 slices, 5 mice | F(2,18) = 2.48<br>p = 0.12 | |
| *b** | Normal distribution | One sample Student's *t* test | 11 cells, 11 slices, 7 mice | p = 0.03 | Discr.: 4.7 ± 1.8, 95% CI: 0.7 to 8.7 |

* Correspond to *a* and *b* in the text, indicating statistical tests that are only mentioned in the text.
